# Capturing large genomic contexts for accurately predicting enhancer-promoter interactions

**DOI:** 10.1101/2021.09.04.458817

**Authors:** Ken Chen, Huiying Zhao, Yuedong Yang

**Affiliations:** School of Computer Science and Engineering, Sun Yat-sen University, 510000, Guangzhou, China; Sun Yat-sen Memorial Hospital, Sun Yat-sen University, 510000, Guangzhou, China; Key Laboratory of Machine Intelligence and Advanced Computing (Sun Yat-sen University), Ministry of Education, Country

**Keywords:** enhancer-promoter interaction, chromatin structure, Transformer, non-coding mutation

## Abstract

Accurately identifying enhancer-promoter interactions (EPIs) is challenging because enhancers usually act on the promoters of distant target genes. Although a variety of machine learning and deep learning models have been developed, many of them are not designed to or could not be well applied to predict EPIs in cell types different from the training data. In this study, we develop the TransEPI model for EPI prediction based on datasets derived from Hi-C and ChIA-PET data. TransEPI compiles genomic features from large intervals harboring the enhancer-promoter pair and adopts a Transformer-based architecture to capture the long-range dependencies. Thus, TransEPI could achieve more accurate prediction by addressing the impact of other genomic loci that may competitively interact with the enhancer-promoter pair. We evaluate TransEPI in a challenging scenario, where the independent test samples are predicted by models trained on the data from different cell types and chromosomes. TransEPI robustly predicts cross-cell-type EPI prediction by achieving comparable performance in cross-validation and independent test. More importantly, TransEPI significantly outperforms the state-of-the-art EPI models on the independent test datasets, with the Area Under Precision-Recall Curve (auPRC) score increasing by 48.84 % on average. Hence, TransEPI is applicable for accurate EPI prediction in cell types without chromatin structure data. Moreover, we find the TransEPI framework could also be extended to identify the target gene of non-coding mutations, which may facilitate studying pathogenic non-coding mutations.

## Introduction

Enhancers are functional DNA fragments acting as cis-regulatory elements on the genome, which regulate gene expressions through chromatin interactions with the promoter of target genes [Maston et al., 2006, Plank and Dean, 2014]. The enhancer-promoter interactions (EPIs) vary highly with cell types and thus play a critical role in cell development and differentiation [Plank and Dean, 2014, Schoenfelder and Fraser, 2019]. EPIs may be disrupted by genetic variations and lead to the dysfunction of genes, underlying the potential pathogenicity of mutations occurring in non-coding regions [Lupiáñez et al., 2015, Li et al., 2018]. Accordingly, linking enhancer mutations to the promoter of target genes could not only help to interpret a substantial number of non-coding mutations but also provide implications for therapeutic approaches [Javierre et al., 2016, Chen and Tian, 2016, Fadason et al., 2018]. However, it remains challenging to identify EPIs accurately because enhancers and their target promoters are typically separated by thousands of base pairs [Schoenfelder and Fraser, 2019, Sanyal et al., 2012].

Previous studies have utilized expression quantitative trait loci (eQTLs) to infer EPIs indirectly [Wang et al., 2013, Wu et al., 2020], while eQTL-based methods are limited to investigate only loci containing variants with high minor allele frequencies due to the requirement of a large number of samples [Consortium, 2015]. Over the last decade, high-throughput chromatin conformation capture-based (3C-based) techniques (e.g., Hi-C [Lieberman-Aiden et al., 2009], ChIA-PET [Fullwood et al., 2009]) have enabled to detect chromatin interactions directly, which could be applied to identify EPIs [Rao et al., 2014, Lu et al., 2020]. However, these 3C-based methods are costly and laborious, experimentally identified EPIs data are available in a few cell types.

To mitigate the problem of identifying EPIs, a variety of computational methods have been proposed to predict EPIs. Though enhancers are sometimes assumed to interact with the nearest promoter, such a method is not reliable because enhancers do not regulate the nearest gene in most cases [Sanyal et al., 2012]. Hence, correlation-based methods were developed later to decipher the underlying rules of EPIs using the correlations of genomic signals at enhancers and genes (or promoters) across a series of cell types [Thurman et al., 2012, Sheffield et al., 2013, Fishilevich et al., 2017]. These methods are of low performance because enhancers are usually cell-type-specific, and EPIs vary across cell types [Moore et al., 2020]. To unveil the complex determinants of EPIs, machine learning (including deep learning) approaches were adopted. Sequence-based methods were developed by making predictions from enhancer and promoter sequences through machine learning or deep learning techniques. Though many methods successfully predict using DNA sequences, such as PEP [Yang et al., 2017], SPEID [Singh et al., 2019], Zhuang’s method [Zhuang et al., 2019], EPIVAN [Hong et al., 2020], the sequence-based methods are inherently cell-type-agnostic and thus are not useful for predicting cell-type-specific EPIs. In parallel, other methods trained machine learning models based on genomic features derived from ChIP-seq and DNase-seq to capture the cell-type-specific characteristics, such as RIPPLE [Roy et al., 2015], TargetFinder [Whalen et al., 2016], JEME [Cao et al., 2017], EPIP [Talukder et al., 2019], EAGLE [Gao and Qian, 2019]. They are more effective than sequence-based models, while they tend to become less inaccurate for predicting EPIs in cell types different from the training data. Besides, the performance of the machine learning models is often exaggerated because the test datasets are not rigorously independent from the training data[Cao and Fullwood, 2019, Moore et al., 2020]. Recently, several methods, like 3DPredictor [Belokopytova et al., 2020] and DeepC [Schwessinger et al., 2020], highlight the importance of employing features of large genomic contexts for chromatin structure modeling. Inspired by these models, we speculate that integrating additional genomic features from large genomic contexts would improve EPI prediction, given that the EPIs are inherently determined by chromatin conformation.

In this study, we present a novel deep learning model named TransEPI for EPI prediction using the Transformer architecture [Vaswani et al., 2017]. TransEPI directly takes the input of genomic signals from large genomic contexts harboring the enhancer-promoter pairs to predict the EPIs. We expected that the features outside enhancers and promoters could enable the model to address the impact of other genomic loci that may competitively interact with the enhancers or the promoters. Given the success of Transformer in protein structure modelling[Jumper et al., 2021, Baek et al., 2021], we adopt a Transformer-based framework in TransEPI for capturing the long-range dependencies between enhancers and promoters. To the best of our knowledge, it is the first model that applies Transformer to predict chromatin interactions. To make an EPI model robust for cross-cell-type prediction, we developed and evaluated TransEPI in a challenging scenario. The TransEPI is trained by a chromosome-split cross-validation scheme, where the training data were split by chromosomes. Then, we tested it on samples where the cell type and chromosome are both different from the training data, as we expected the evaluation could reliably reflect the actual performance of models. Our TransEPI method is shown robust by achieving comparable performance in cross-validation and the independent test datasets. More importantly, TransEPI significantly outperforms two state-of-the-art EPI models, with an average auPRC increase by 48.84 % on the test datasets. Additionally, we found our TransEPI framework is also helpful for predicting cell-type-specific target genes affected by non-coding mutations.

The implementation of TransEPI and the datasets are available at https://github.com/biomed-AI/TransEPI.

## Materials and Methods

### Datasets

We employed the BENGI dataset to develop TransEPI for predicting EPIs and the Hi-C loop dataset to extend TransEPI for identifying the target gene of non-coding mutations.

*The BENGI dataset* We recruit data from the “Benchmark of candidate Enhancer-Gene Interactions (BENGI)” dataset [Moore et al., 2020] to develop the TransEPI model. BENGI is a collection of enhancer-target gene pairs from several cell lines or tissues identified by 3C-based methods or genetic approaches. Since we aimed to predict the physical interactions between enhancers and promoters, only the samples identified by Hi-C (GM12878, HeLa-S3, HMEC, IMR90, K562, and NHEK) and ChIA-PET (GM12878 and HeLa-S3) were utilized.

Because the samples curated by BENGI are enhancer-gene interactions, we first mapped the genes to transcripts based on GENCODE annotation (GRCh37/hg19) [Frankish et al., 2019]. Then, by defining the 1500-base pair (bp) upstream and the 500-bp downstream of transcript start site (TSS) as the promoter, we obtained the enhancer-promoter(EP) pairs we need for developing our TransEPI model. Since even the TSSs of the same gene may be thousands of base pairs apart, the EP pairs derived from positive samples but with promoters residing outside the anchor region of chromatin loops were discarded. Besides, we also removed the samples with low-expressed transcripts (transcript per million (TPM) < 1) from both positive and negative samples as they were less likely to be regulated by enhancers (We did this to eliminate false-positive samples as many as possible). Finally, we obtained 45,182 positive and 307,135 negative samples (Table 1) from 6 cell lines.

**Table 1.**
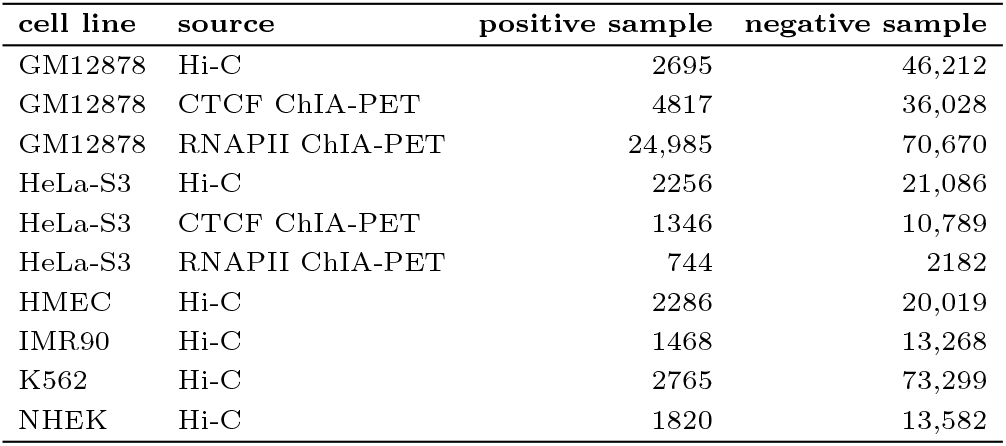
Summary of the dataset

| cell line | source | positive sample | negative sample |
| --- | --- | --- | --- |
| GM12878 | Hi-C | 2695 | 46,212 |
| GM12878 | CTCF ChIA-PET | 4817 | 36,028 |
| GM12878 | RNAPII ChIA-PET | 24,985 | 70,670 |
| HeLa-S3 | Hi-C | 2256 | 21,086 |
| HeLa-S3 | CTCF ChIA-PET | 1346 | 10,789 |
| HeLa-S3 | RNAPII ChIA-PET | 744 | 2182 |
| HMEC | Hi-C | 2286 | 20,019 |
| IMR90 | Hi-C | 1468 | 13,268 |
| K562 | Hi-C | 2765 | 73,299 |
| NHEK | Hi-C | 1820 | 13,582 |

We combined the samples from GM12878 and HeLa-S3 to construct a training set, namely BENGI-train, which contains 36,843 positive and 186,967 negative ones. The samples from the other 4 cell lines are all reserved for independent test, namely BENGI-HMEC, BENGI-IMR90, BENGI-K562, and BENGI-NHEK, respectively.

*The HiC-loop dataset* Because most of the non-coding mutations reside outside the putative enhancer regions, the original TransEPI model trained on BENGI fails to help find target genes of mutations. Therefore, we extended the TransEPI model for predicting general chromatin interactions by training it on a novel dataset consisted of Hi-C loops. To this end, we obtained Hi-C loops in 7 cell lines (GM12878, HeLa-S3, HMEC, HUVEC, IMR90, K562, and NHEK) from the Gene Expression Omnibus (GEO) database under the accession number of GSE63525 [Rao et al., 2014].

Positive samples are defined as the pairs of Hi-C loop anchors. The negative samples were generated by randomly pairing the anchor regions of loops and the other randomly selected regions from the genome. Notably, the distances between the loop anchors are not in the uniform distribution (Figure S2). Therefore, we intentionally sampled more samples (about 50 %) matching the E-P distribution of positive samples to avoid the model predicting EPIs by simply using the E-P distance.

Finally, we compiled the Hi-C loop dataset consisting of 38, 608 positive and 272, 397 negative samples (Details shown in Table S2)

### Input features

TransEPI is designed to take the input of features from large genomic regions harboring the candidate E-P pairs (Figure 1a). We set the length of large genomic regions to 2,500,000 bp because the maximum value of E-P distance in BENGI is 2, 246, 878 and there are only 30 samples with E-P distance longer than 2, 000, 000 bp. The start and the end of the large genomic regions are determined by extending 1,250,000 bp from the midpoint of the E-P pair up- and down-stream. When the enhancer or promoter is close to the ends of chromosomes, we will shift the region to keep the region within the range of chromosomes.

**Fig. 1.**
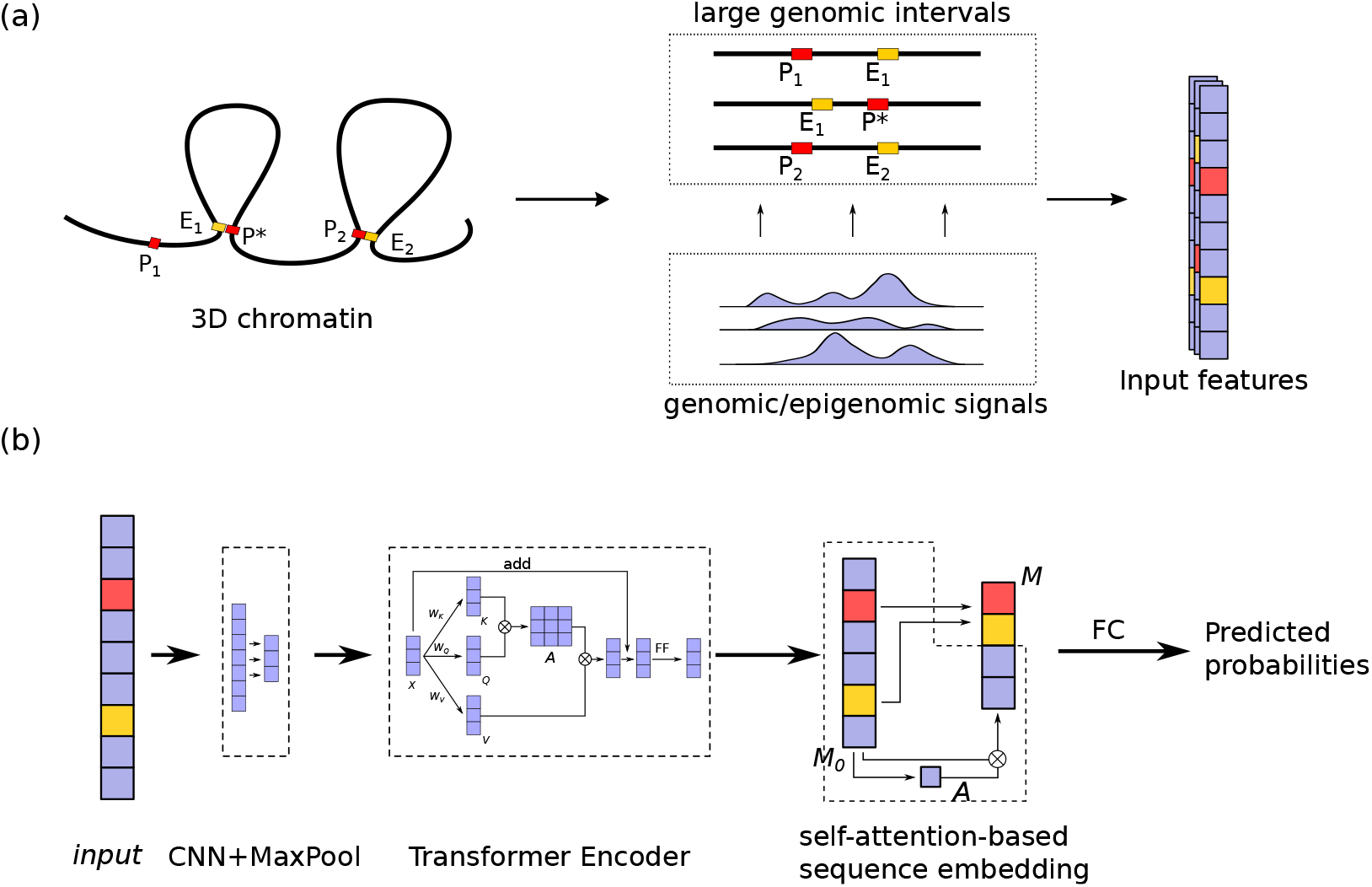
The TransEPI framework. (a) Feature preparation. Genomic features (CTCF, DNase I, H3K27me3, H3K4me1, H3K4me3, H3K36me3, and H3K9me1) are extracted from large intervals harboring the candidate enhancer-promoter pairs (enhancers are marked in yellow and promoters are marked in red); (b) The architecture of the TransEPI. TransEPI are mainly consisted of 3 modules, a CNN+MaxPooling module extracting features from the input sequences, a Transformer Encoder module capturing the long-range dependencies between enhancers and promoters, and a self-attention-based sequence embedding module encoding the sequential features into a low-dimensional embedding. Finally, the embedding of the whole sequence and the hidden states of the enhancer bin and the promoter bin are concatenated together to predict the probability with a fully-connected layer. (⊗: matrix multiplying operation: CNN: convolutional neural network; MaxPool: max-pooling; FC: fully-connected)

Inspired by Belokopytova et al[Belokopytova et al., 2020], we partitioned each 2.5Mbp region into 5000 consecutive bins using a bin size of 500 bp and averaged the genomic and epigenomic signals within each bin to represent the chromatin states. Here, the genomic and epigenomic signals include CTCF binding sites, chromatin accessibility (DNase-I signals), and 5 histone modification marks (H3K4me1, H3K4me3, H3K27me3, H3K36me3, and H3K9me3). The CTCF binding sites in narrowPeak format for each cell line were obtained from the Encyclopedia of DNA Elements (ENCODE) project [Davis et al., 2018]. The DNase-I and histone marks data in bigWig format were taken from the Roadmap Epigenomics Project [Kundaje et al., 2015]. Apart from the genomic features, we also encoded the relative distance to the enhancer or the promoter for each bin as an additional feature to make the model aware of the locations of the E-P. Details about feature preparation are described in Supplementary Methods.

### The TransEPI model

The architecture of the TransEPI model is illustrated in Figure 1b.

Firstly, TransEPI utilizes a one-dimensional convolutional neural network (1D CNN) to exact features from the input signals. A max-pooling layer is then used to down-sample the features and shrink the length of each input sequence (The output sequence will be denoted by ***X*** = [*x*_1_, *x*_2_, · · · , *x*_*l*_], where 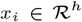, *h* is the dimension of the hidden states).

Next, we employ the Transformer encoder module to capture the long-range dependencies between enhancers and promoters within the sequential signals ***X***. ***X*** is first transformed into a key, a query, and a value sequence, respectively:

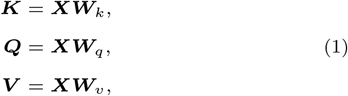

where 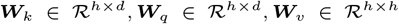 are learnable weight matrices. ***K*** and ***Q*** are then used to construct the attention matrix ***A***:

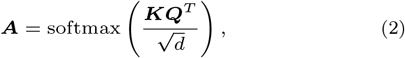

where the attention coefficient *a*_*i,j*_ in ***A*** could be understood as the correlation between the *i*-th and the *j*-th position in ***X***. By multiplying ***V*** by ***A***, the hidden states at different positions are exchanged and updated based on ***A***. To make the Transformer model deeper, multiple Transformer encoder layers could be stacked sequentially. The output from the Transformer module is denoted by 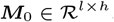.

Thereafter, we employ a self-attention-based sequence embedding module [Lin et al., 2017] to obtain a low-dimensional embedding for each sequence. Specifically, we feed ***M***_0_ into a two-layer fully-connected (FC) network to obtain the attention coefficients (weights) for different positions in ***M***_0_ and then multiply ***M***_0_ by the weights to obtain a weighted embedding 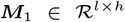. Then, we apply the average and the maximum pooling over all the hidden states in ***M***_1_ and concatenate them with the hidden states corresponding to the location of the enhancer (***h**_e_*) and the promoter (***h***_*p*_):

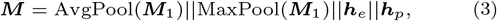

where 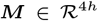. It includes the global features from the whole sequence and the local features from the enhancer and the promoter.

Finally, the final sequence embedding ***M*** is passed to a two-layer FC module to predict the EPI:

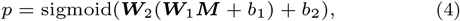

where 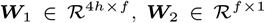, *b*_1_, *b*_2_ are all the weights in the FC module. As a Sigmoid function is used, *p* ranges from 0 to 1 (*p* ∈ (0, 1), representing the probability that the input enhancer and the promoter interact with each other. Meanwhile, in order to make TransEPI sensitive to the location of the enhancer and the promoter, we use another FC module to predict the E-P distance:

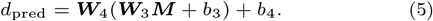

### Evaluation metrics

We evaluated the model using the Area Under the Precision-Recall (PR) curve (auPRC) and the Area Under the Receiver Operating Characteristic (ROC) curve [Hanley and McNeil, 1982] (AUC). The PR curve is a plot of precision against the recall at a series of thresholds. Similarly, the ROC curve is a plot of true-positive rate (TPR) against the false-positive curve (FPR). Precision, recall, TPR, and FPR are defined as:

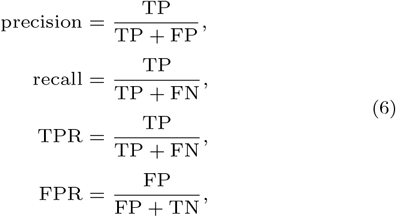

where TP, FP, TN, and FN are short for the True Positives (correctly predicted interacting pairs), False Positives (falsely predicted non-interacting pairs), True Negatives (correctly predicted non-interacting pairs), and False Negatives (falsely predicted interacting pairs).

Since the auPRC is associated with the ratio of the number of positive and negative samples, we also used auPRC-ratio (dividing auPRC by the proportion of positive samples [Pratapa et al., 2020]) as a metric for comparing performance across datasets.

### Model training and evaluation

The TransEPI model is implemented with PyTorch (version 1.9.0) [Paszke et al., 2019] in Python 3.8. It was trained to minimize the binary cross entropy loss for EPI prediction and the mean squared error (MSE) loss for E-P distance prediction simultaneously:

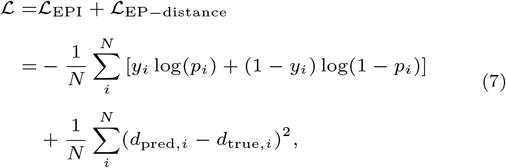

where *p_i_*, *y_i_*, *d*_pred*,i*_, and *d*_true*,i*_ are the the predicted EPI probability, true EPI label (0 or 1), the predicted E-P distance, the true E-P distance of the *i*-th sample, respectively. We used the Adam optimizer [Kingma and Ba, 2017] to update the weights in the neural network. The early stopping strategy was utilized for regularization.

In order to avoid over-fitting, we adopt a cross-chromosome 5-fold cross-validation scheme to fine-tune the hyper-parameters (Figure S1) in TransEPI. We divided the samples in BENGI-train into 5 folds by chromosomes, ensuring that the samples from the same chromosome would also be put into the same fold (chromosomes assigned to each fold are listed in Table S1). It is a critical approach to avoid over-fitting for building machine learning models in genomics [Schreiber et al., 2020]. In each training epoch, we trained the model on 4 folds and validated it on the remaining fold in turn. The average AUC and AUPR on the 5 folds are used to measure the performance. For evaluating our method in a challenging way, the independent test sets are also split into 5 folds by chromosomes according to the partition table in Table S1. The test samples are only predicted by the models trained on different chromosomes, as illustrated in Figure S1.

### State-of-the-art methods for comparison

We compared TransEPI to two state-of-the-art methods: TargetFinder and 3DPredictor.

#### TargetFinder

The TargetFinder model is a Gradient Boosting model using various genomic features [Whalen et al., 2016]. As shown in two studies [Xi and Beer, 2018, Cao and Fullwood, 2019], the reported performance in the original paper was inflated because the test data was not rigorously independent from the training data. However, it still outperforms many other EPI models, according to a benchmark study by Moore et al [Moore et al., 2020]. Thus, we used TargetFinder as the first baseline model for comparison. Given that no official implementation of TargetFinder is available, we implemented it in Python using the XGBoost package, which is a state-of-the-art gradient boosting model [Chen and Guestrin, 2016]. The genomic features used in TargetFinder are the same as our TransEPI model.

#### 3DPredictor

The 3DPredictor model [Belokopytova et al., 2020] is an XGBoost model which was originally developed to identify enhancer-promoter contacts via predicting the Hi-C contact map. It demonstrates that integrates only integrating the oriented CTCF binding peaks within and around the pair of chromatin loci could achieve accurate predictions. Here, we adapted it to directly predict EPIs.

For a fair comparison, the baseline models and TransEPI were trained on the same datasets. The hyper-parameters in the baseline models are rigorously refined using the random grid search strategy.

The source codes associated with data preparation and model training of the baseline models are available in our Github repository, as well.

### Identifying target genes for non-coding mutations

We extend the TransEPI framework to predict the target gene of non-coding mutations. As the original TransEPI is built for predicting EPIs, it is not directly applicable to predict mutation-gene pairs because we expect it to be applicable for all non-coding mutations, even the mutations residing outside enhancer regions. Hence, we trained TransEPI on the Hi-C loop dataset, using HMEC, HUVEC, and the remaining cell lines as the validation, test, and training datasets, respectively.

For identifying the target genes, a candidate mutation-transcript list is first curated for each mutation by pairing it with the transcripts (transcript’s TSS) within 1M bp. Subsequently, TransEPI is applied on the mutation-TSS pairs using genomic features from two neural cells and 3 brain tissues (Table S10). The mutation-TSS pair with a predicted probability above a certain threshold will be kept as interacting pairs.

## Results

### TransEPI accurately predicts EPIs in different cell types

EPIs are cell-type-specific chromatin interactions that vary significantly across cell types. To make TransEPI applicable for cross-cell-type EPI prediction, it is critical to avoid over-fitting when we develop and evaluate the model [Schreiber et al., 2020]. To this end, we not only adopt the chromosome-split 5-fold cross-validation scheme to train and fine-tune the model but also test it on samples from different cell types and chromosomes (Figure S1).

We first compared the results of TransEPI on cross-validation and independent test datasets to investigate whether it could achieve consistent performance across cell types. As shown in Figure 2a, the auPRC-ratios achieved by TransEPI on 4 independent test datasets are 6.516, 5.847, 5.301, and 3.717, 3 out of which are higher (in red) than that on the cross-validation dataset (auPRC-ratio=4.278). Moreover, such a trend is consistently observed in the results on each fold. As such, the results suggest that TransEPI trained on BENGI-train can be well applied to new cell types.

**Fig. 2.**
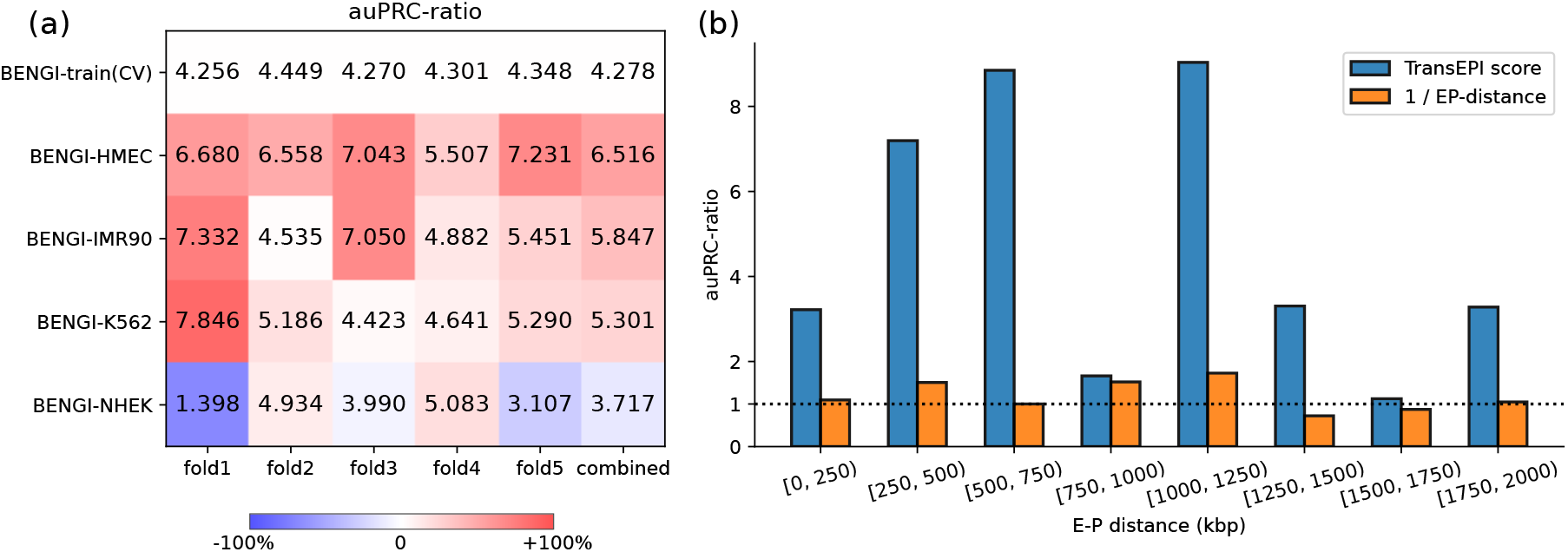
Evaluating TransEPI on independent test datasets. (a) The auPRC-ratio scores of TransEPI on BENGI-train(CV) and 4 independent tests. (b) The auPRC-ratio scores of TransEPI on independent test samples stratified by distance. TransEPI consistently outperforms the naive distance-based method in every group.

Previous studies have found that the distance from enhancer to the promoter (EP-distance) may have a strong predictive power on some datasets [Gao and Qian, 2019, Moore et al., 2020]. However, in fact, the predictive power of the EP-distance is determined by the distribution of EP-distance in negative samples, which could be controlled by the way of negative sampling. In order to eliminate potential bias caused by EP-distance, we additionally evaluated TransEPI on test datasets stratified by EP-distance. Specifically, we merged the samples in BENGI-HMEC, BENGI-IMR90, BENGI-K562, and BENGI-NHEK and stratified the samples by E-P distance into 8 groups using the bin size of 250,000 bp. In this way, we ensured that using only EP-distance can hardly predict EPIs in each group, as the auPRC-ratio ranges from 0.7222 to 1.727 (a random model is expected to achieve the auPRC-ratio of 1). In contrast, TransEPI achieves much higher auPRC-ratios, which range from 1.125 to 9.036, on the E-P distance-stratified datasets (Figure 2b), demonstrating that TransEPI does capture the underlying determinants of EPI instead of just relying on E-P distance alone.

Taken together, we could conclude that TransEPI is capable of deciphering the mechanisms of enhancer-promoter interaction that are universal to different cell types.

### TransEPI significantly outperforms the baseline methods

To further evaluate TransEPI, we compared it with two state-of-the-art methods, TargetFinder [Whalen et al., 2016] and 3DPredictor [Belokopytova et al., 2020], on the independent test datasets.

In Figure 3a, the ROC plots show that TransEPI significantly outperforms TargetFinder and 3DPredictor on BENGI-HMEC and BENGI-IMR90 (all the *P*-values < 1 × 10^−5^, by McNeil & Hanley’s test on AUC, as listed in Table S3). On BENGI-K562 and BENGI-NHEK, although the AUC of TransEPI does not surpass that of TargetFinder and 3DPredictor, TransEPI tends to have a higher TPR when the FPR scores are close to 0. This means that TransEPI can find more interacting E-P pairs when controlling false positive rate at a low level, implying that TransEPI is more helpful for practical use. When we compare these models by auPRC (Figure 3b), TransEPI consistently outperforms TargetFinder and 3DPredictor on all the test datasets. The average auPRC of TransEPI increases by 48.84 % compared with the second best method TargetFinder, further demonstrating the superior performance of TransEPI to the other methods.

**Fig. 3.**
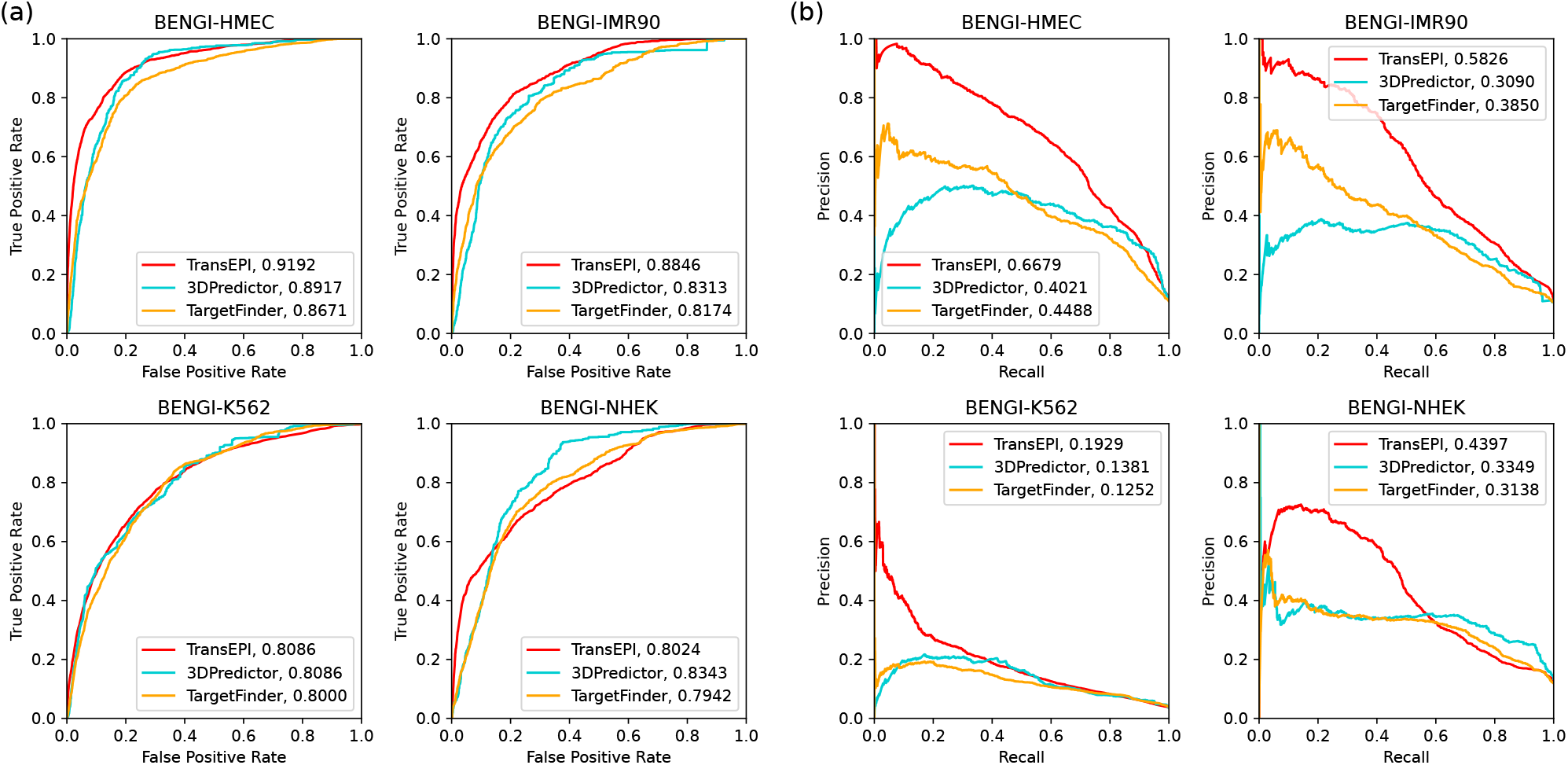
Comparing TransEPI with TargetFinder and 3DPredictor on 4 independent test sets. (a) Receiver Operating Characteristic (ROC) curves and (b) Precision-recall (PR) curves are plotted.

Compared to our TransEPI model, TargetFinder employs only the average genomic signals within the genomic windows between enhancers and promoters. It does not only lack a fine representation of the features in genomic windows but also completely ignores the features within the outer regions of the enhancer-promoter pairs. Therefore, we believe that it is important to use a finer features representation strategy and leverage features from larger genomic contexts.

As for 3DPredictor, its authors employ only the CTCF binding site as a feature for chromatin structure modelling as they found additional features could not provide a significant performance improvement. However, in our study, additional features are found beneficial (See Supplementary Methods and Table S4). We speculate that this is because the EPIs may have distinct characteristics from the common chromatin interactions, as the activity of promoters and enhancers are usually inferred from chromatin accessibility and histone modification marks [Tsompana and Buck, 2014, Gao and Qian, 2020].

### TransEPI benefits from the features outside enhancers and promoters

In this section, we quantitatively studied the contribution of features from the window regions between the enhancers and promoters (“window features”) and the neighboring regions outside the enhancer-promoter pairs (“neighbor features”). To this end, we masked the window or the neighbor features by setting the genomic features within the window (the w/o W setting) or the neighbor (the w/o N setting) regions to 0, respectively. In both settings, the features in the enhancer and promoter bin along with the 2 bins up- and down-stream of them are kept. Additionally, we also conducted a w/o NW setting by masking both the window and the neighbor features.

As shown in Figure 4a, masking window and neighbor features decrease the auPRC scores on all 4 independent test datasets. The exclusion of window features has the most prominent effect, with the average auPRC score decreasing by 41.45 % (Table S5). Such an observation is in concordance with previous studies [Whalen et al., 2016, Gao and Qian, 2019], in which they demonstrate the importance of window features. More importantly, we found that the neighbor features are non-trivial, as well. The absence of neighbor features decreases the average auPRC score by 13.08 % and masking both neighbor and window features results in the average auPRC score decreasing by 54.96 %, on the 4 independent test datasets (Table S5).

**Fig. 4.**
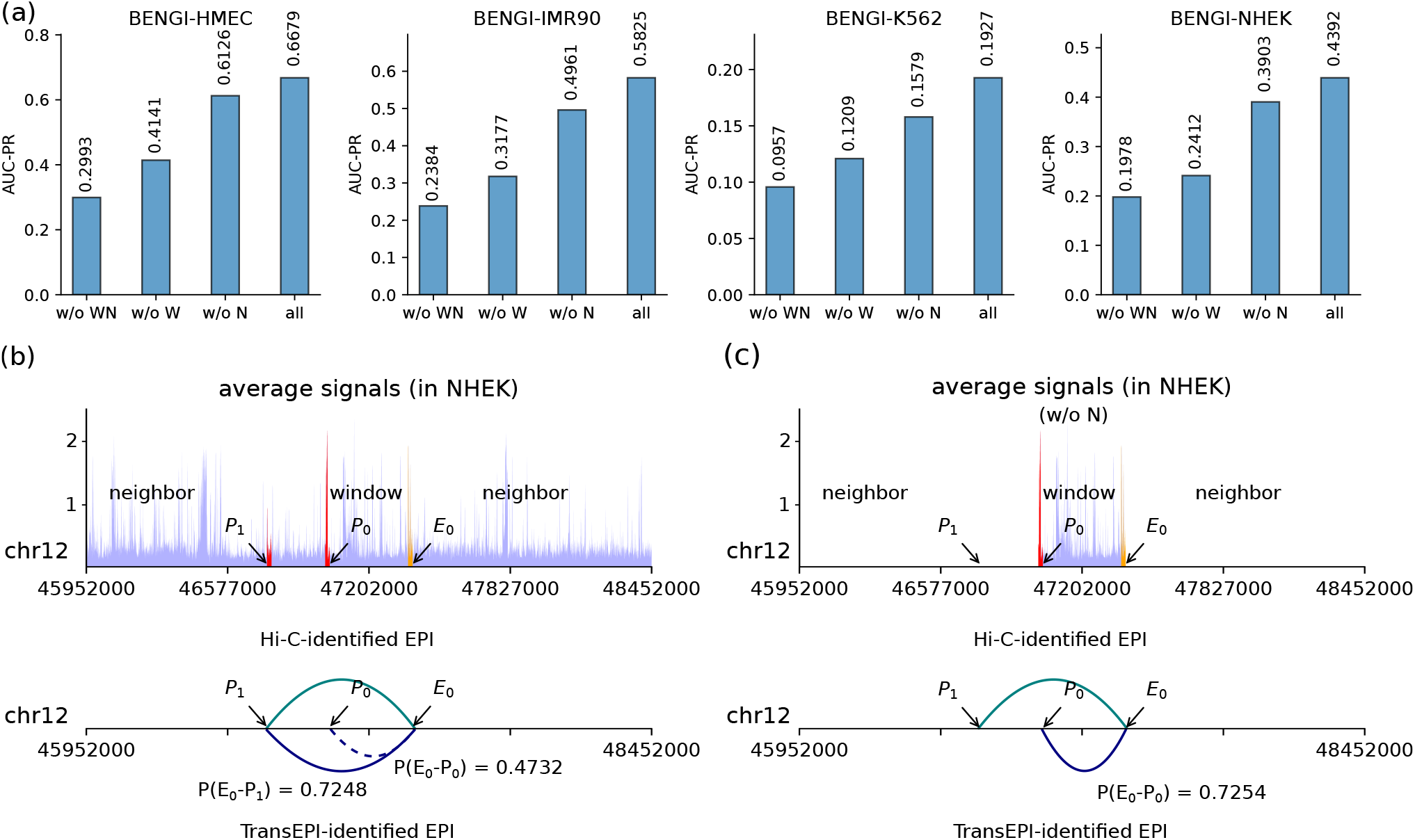
Ablation study on features outside enhancers and promoters. (a) We reported the average auPRC of TransEPI model using all the features (all), masking features in the neighbor regions (w/o N), masking features in window regions (w/o W), and masking feature in both neighbor and window regions (w/o NW) on 4 independent test sets, where the error bar stands for the standard deviation; (b) E_0_ (EH37E0265448) is an enhancer in the NHEK cell line. P_0_ and P_1_ are promoters of transcripts ENST00000548870 and ENSG00000139211, respectively. When we use both the neighbor and the window features, TransEPI correctly predicts E_0_-P0 and E_0_-P_1_ as non-interacting (marked with dotted line) and interacting pairs (marked with solid line), respectively. (c) When we mask the neighbor features, TransEPI mistakenly identifies E_0_-P_0_ as an interacting pair. The genomic signals shown in (b) and (c) are the average over different features.

A case study under the w/o N setting intuitively explains why the features outside enhancers and promoters are indispensable. In cell line NHEK, the enhancer *E*_0_ (EH37E0265448,chr12:47,070,581-47,071,150) interacts with the promoter *P*_1_ (ENSEMBL transcript ID: ENST00000548870, TSS=chr12:46,761,193), but not the promoter *P*_0_ (ENSEMBL transcript ID: ENST00000550413, TSS=chr12:47,473,425). The promoter *P*_1_ resides at the neighboring region of *E*_0_-*P*_1_. When we use all the features, TransEPI correctly identifies *E*_0_-*P*_1_ as an interacting pair (*P* (*E*_0_-*P*_1_) = 0.7248) and *E*_0_-*P*_0_ as a non-interacting pair (*P* (*E*_0_-*P*_0_) = 0.4732), respectively (taking 0.5 as the threshold) (Figure 4b). When we mask the neighbor features, TransEPI is not able to aware of the existence of promoter *P*_0_. As a result, TransEPI mistakenly regards *E*_0_-*P*_0_ as an interacting pair, with *P* (*E*_0_-*P*_0_) increasing from 0.4732 to 0.7254 (Figure 4c). The case study suggests the other regulatory elements (elg.: enhancers, promoters) around the enhancer-promoter pair may competitively interact with them. Considering only the chromatin states of the enhancer-promoter pair is far from enough for predicting EPIs accurately.

### TransEPI facilitates identifying target genes of non-coding mutations

Explaining GWAS results may be a challenging task because a large number of the risk variants reside in non-coding regions whose functions are not well characterized. An effective solution is to link the variants to target genes and previous studies have successfully employed Hi-C or eQTL data for explaining GWAS results [Sey et al., 2020, Chen and Tian, 2016, Lu et al., 2020]. However, the Hi-C and eQTL data are tissue-specific and may not be available in the tissue or the cell line studied by researchers. Here, given the outstanding performance of TransEPI for predicting EPIs, we extended to predict tissue-specific target genes of non-coding variants.

We collected the mutations associated with neural diseases or brain disorders sorted by Lu et al [Lu et al., 2020], which are taken from the GWAS Catalog [Buniello et al., 2019]. Only the non-coding mutations residing in intergenic or intronic regions are kept, by which we obtained 3943 non-coding mutations (Table S6). Using TransEPI, we identified 5131 mutation-target gene pairs associated with 3034 genes and 2571 mutations (Table S7), only protein coding genes are included). We firstly conducted Gene Ontology (GO) analysis on the target genes using gProfiler [Raudvere et al., 2019]. As shown in Table S8, the target genes significantly enrich 400 GO terms, which suggests that TransEPI-predicted genes are of biological significance. More importantly, we found various neural function-associated GO terms and 5 out of the top-10 GO biological process (GO:MF) are relevant to neural functions (Figure 5a). As a case study, TransEPI correctly identifies the target gene of two mutations: rs10153620 (NRP2, TransEPI-score = 0.9100) [Ebejer et al., 2013] and rs10457592 (POU3F2, TransEPI-score = 0.9400)[Hyde et al., 2016], which have been validated using Hi-C[Lu et al., 2020].

**Fig. 5.**
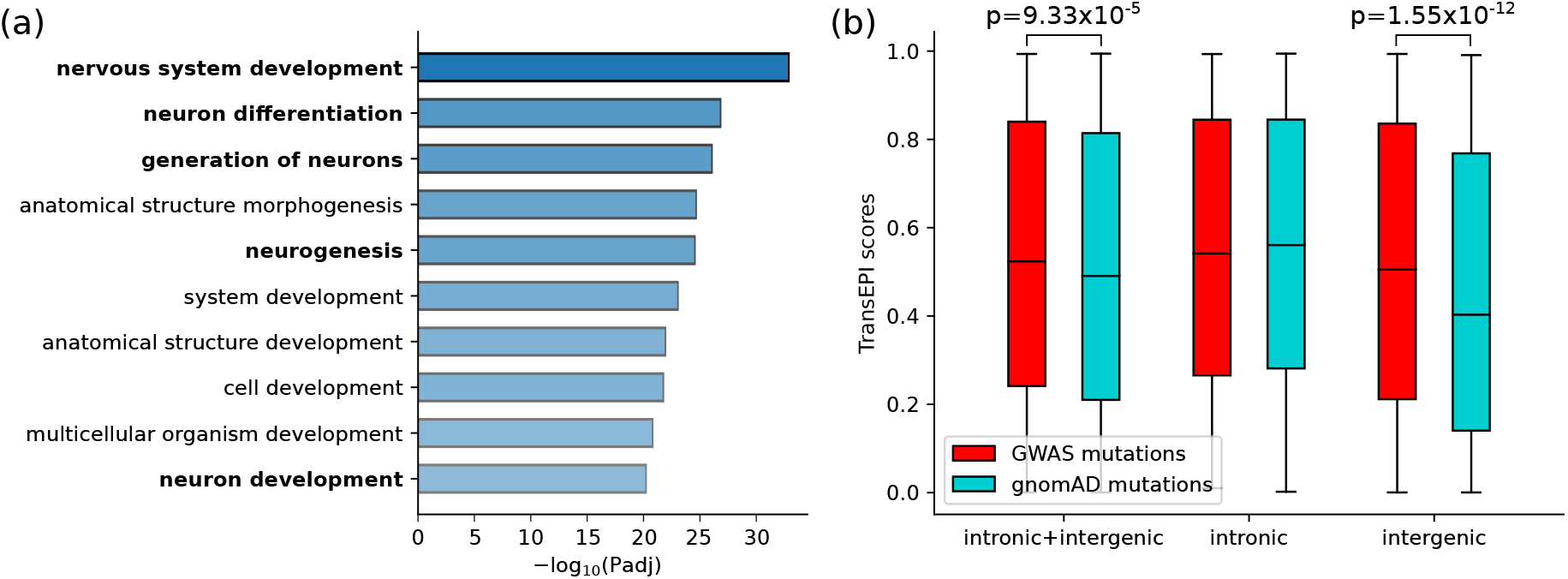
Applying TransEPI to find target genes for non-coding mutations. (a) The top-10 Gene Ontology Biological Process (GO:BP) items enriched by TransEPI-predicted protein-coding genes for a group of non-coding variants related to neural diseases or brain disorders. There are 5 GO items associated with neural functions among the top-10 GO items. (b) Comparison of the TransEPI-predicted top-1 scores of GWAS mutations and gnomAD mutations in the non-coding (intronic+intergenic), the intronic, and the intergenic group. Statistical significance is assessed by *t*-test.

The above analysis implies that TransEPI-predicted genes may be functionally associated with neural functions. However, the statistical significance of our observations can not be assessed since we lack the ground-truth target genes for most mutations. To further evaluate the predictions, we adopted an alternative approach by comparing the predicted target genes of disease-related mutations to those of disease-irrelevant mutations: We first extracted the highest TransEPI-predicted probability, namely top-1 score, among all the candidate target genes for each mutation. The top-1 score represents the probability that a mutation interacts with at least one target gene. We believe that the disease-related mutations are expected to be more likely to interact with target genes than the disease-irrelevant ones. Accordingly, the top-1 scores of the disease-related mutations should be higher than those of the disease-irrelevant ones. Specifically, we used the above-mentioned GWAS mutations as disease-related mutations (GWAS-mutations) and compiled a group of disease-irrelevant non-coding mutations (n = 19,715, 5 times that of GWAS mutations) from the gnomAD database (gnomAD-mutations) (Table S9). As shown in Figure 5b, the disease-related mutations have significantly higher top-1 scores than the disease-irrelevant ones (P-value = 9.33 × 10^−5^, by *t*-test). Next, we split non-coding mutations into an intronic and an intergenic group and compared the predictions for them, respectively. We observed a more significant difference in the intergenic group (P-value=1.55 × 10^−12^), while no significant difference is found in the intronic group. This is likely because that the intergenic mutations are more likely to affect distal target genes than intronic mutations.

Taken together, we could conclude that the TransEPI framework is also helpful to identify the target gene of non-coding mutations and thus could potentially facilitate explaining GWAS results.

## Discussion

In this study, we present a novel deep learning model, TransEPI, for predicting EPIs by capturing large genomic contexts using the Transformer architecture. Instead of considering only the states of an individual pair of enhancer and promoter (EP), TransEPI takes the whole environment where they locate into account. In this way, TransEPI could be aware of the impact of other regulatory elements that may competitively interact with the EP and hence achieve the state-of-the-art performance for EPI prediction.

A variety of EPI models have been proposed yet, while many of them suffer from exaggerated performance and are not well applicable for cross-cell-type EPI prediction [Xi and Beer, 2018, Cao and Fullwood, 2019, Moore et al., 2020]. This is because the datasets used for training and validation (or test) are randomly separated. Therefore, samples with sharing features may be included in both the training and the validation data, resulting in severe over-fitting caused by data leakage. To alleviate the problem of over-fitting, we train and fine-tune the TransEPI model through 5-fold cross-validation (CV), where the data are split by chromosomes to ensure that the samples in different folds do not overlap with each other. Besides, we evaluate TransEPI on independent datasets derived from 4 cell lines to assess whether it could predict EPIs in cell lines different from the training data. As TransEPI achieves comparable performance in CV and on independent test datasets, the chromosome-split cross-validation scheme is shown to be effective to avoid over-fitting.

Since EPIs are inherently determined by the conformation of chromatin, we speculated that additional enhancers and promoters around the E-P pair to be predicted are all critical for accurate EPI prediction. To effectively capture the long-range dependencies between the enhancers and promoters, we present the TransEPI framework which is mainly based on the Transformer encoder architecture. By ablation study, we found TransEPI is sensitive to the additional regulatory elements that may competitively interact with the E-P pair to be predicted, demonstrating the importance of using large genomic contexts in EPI models.

Given that TransEPI enables accurate EPI prediction, we extended the framework to find the target genes of non-coding mutations. By applying the model on mutations associated with neural diseases or brain disorders, TransEPI found target genes that are functionally associated with neural functions. Moreover, these disease-associated non-coding mutations are found to have a higher ratio to act on target genes than those irrelevant to diseases.

Although TransEPI has achieved the state-of-the-art performance, there is still much room for improvement. For example, the time complexity and memory usage required by the standard Transformer module [Vaswani et al., 2017] are quadratic to the length of the input sequence, which are computationally expensive. It is infeasible for us to take larger genomic contexts into consideration in our model. In the future, we may leverage more light-weight Transformer architectures [Zhou et al., 2020, Choromanski et al., 2020, Katharopoulos et al., 2020] to alleviate the problem. As for datasets, we currently leverage only EPIs identified by Hi-C and ChIA-PET. Future versions of TransEPI could consider using EPIs identified by the other 3C-based methods like capture Hi-C [Mifsud et al., 2015] and HiChIP[Mumbach et al., 2016], which are more suitable to detect EPIs. Besides, due to the resolution of 3C-based experiments, the assignment of enhancers to target promoters may be ambiguous and thus make the training data noisy. It is of interest to explore how to integrate additional evidence (e.g.: eQTL mapping) or employ the models enhancing the resolution of Hi-C data [Zhang et al., 2018, Li and Dai, 2020]to improve the quality of training data.

## Acknowledgments

This study has been supported by the National Key R&D Program of China (2020YFB020003), National Natural Science Foundation of China (61772566), Guangdong Key Field R&D Plan (2019B020228001 and 2018B010109006), Introducing Innovative and Entrepreneurial Teams (2016ZT06D211), Guangzhou S&T Research Plan (202007030010).

