## Supplementary Methods for "Capturing large genomic contexts for accurately predicting enhancer-promoter interactions"

### 1 Supplementary Figures

#### 1.1 Illustration of the cross validation and test pipeline

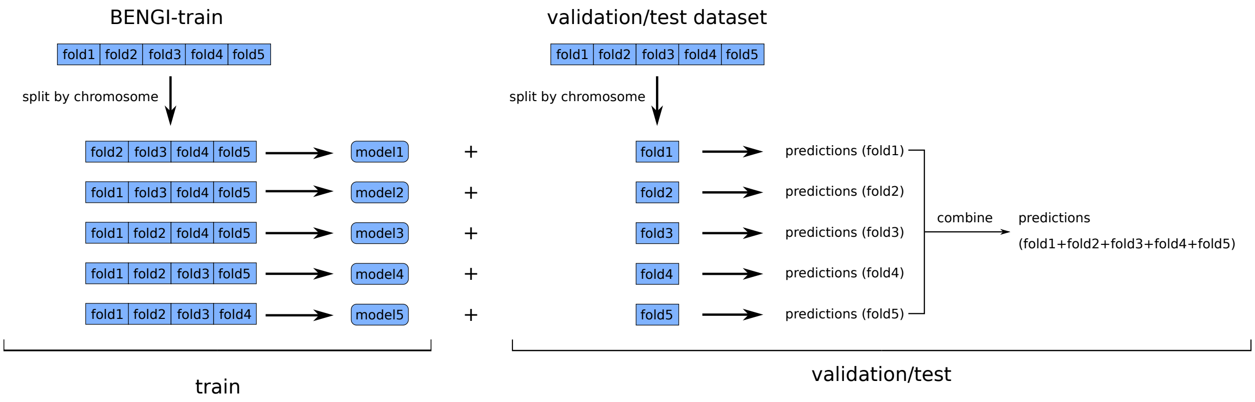

Figure S1: Illustration of the cross validation and test pipeline.
fold1: chr1, chr10, chr15, chr21;
fold2: chr19, chr3, chr4, chr7, chrX;
fold3: chr13, chr17, chr2, chr22, chr9;
fold4: chr12, chr14, chr16, chr18, chr20;
fold5: chr11, chr5, chr6, chr8

#### 1.2 Distribution of Hi-C loop length

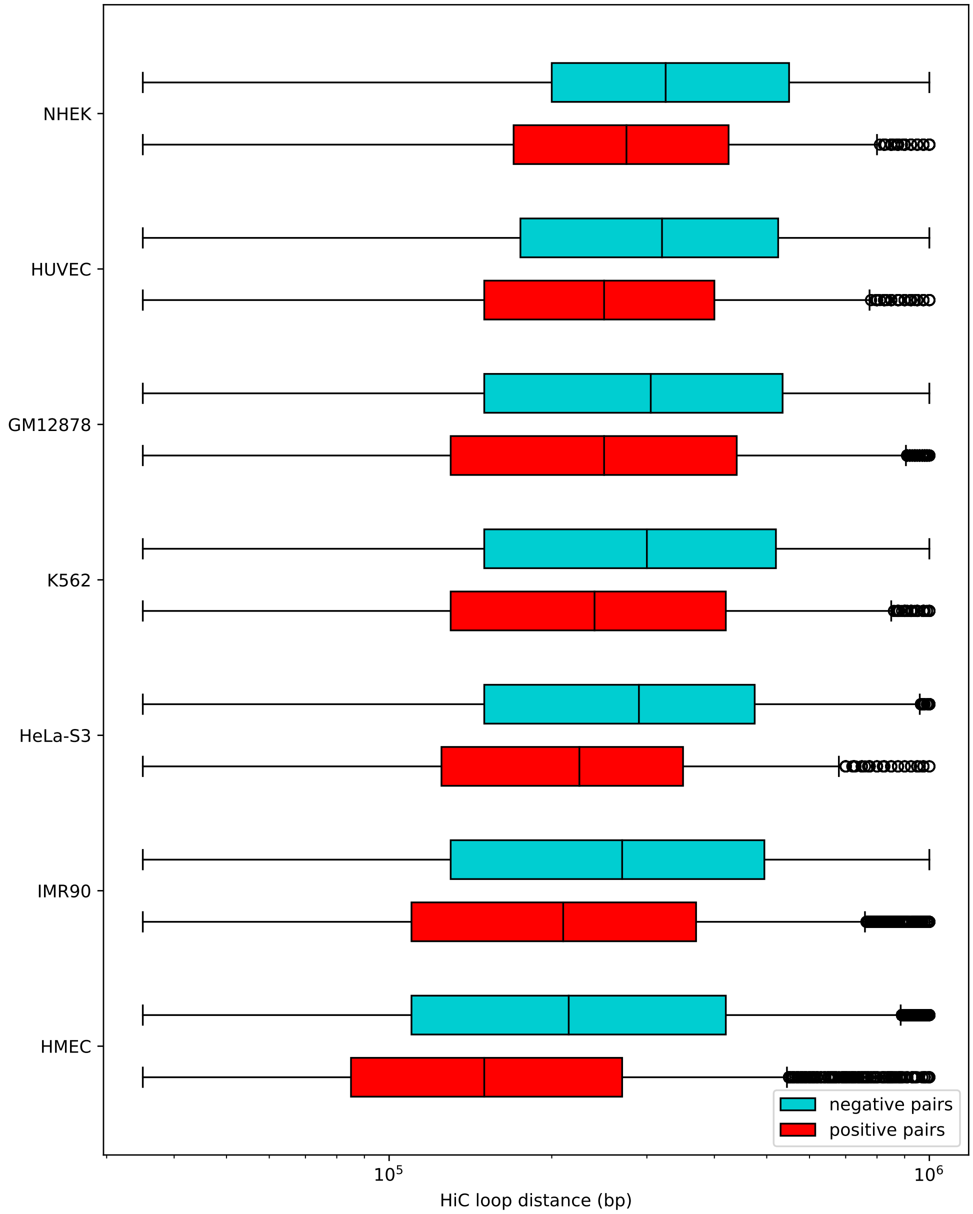

Figure S2: The loop distance distribution in positive and negative samples in the Hi-C loop dataset

#### 1.3 Distribution of EP-distance

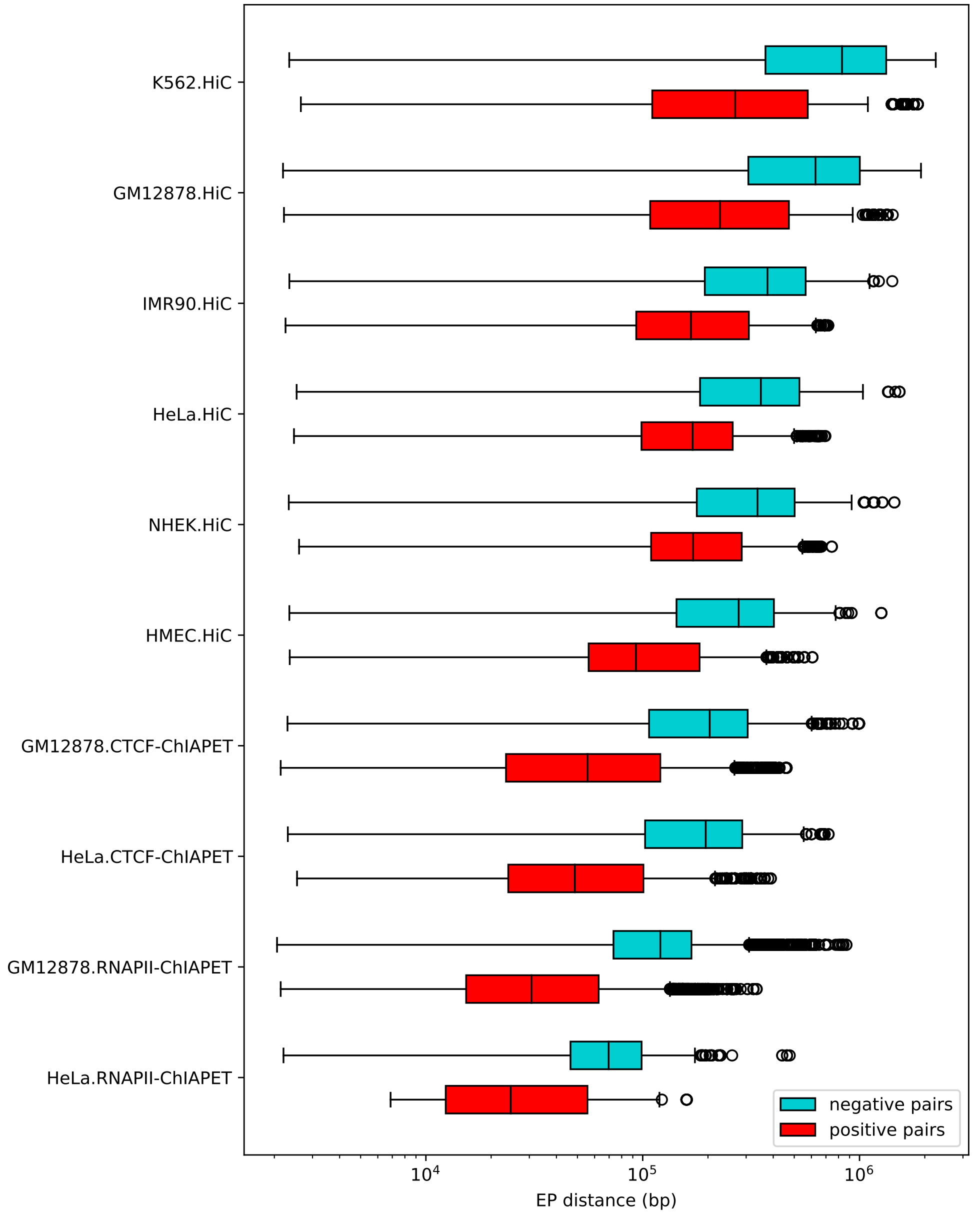

Figure S3: The EP distance distribution in positive and negative samples in the BENGI dataset

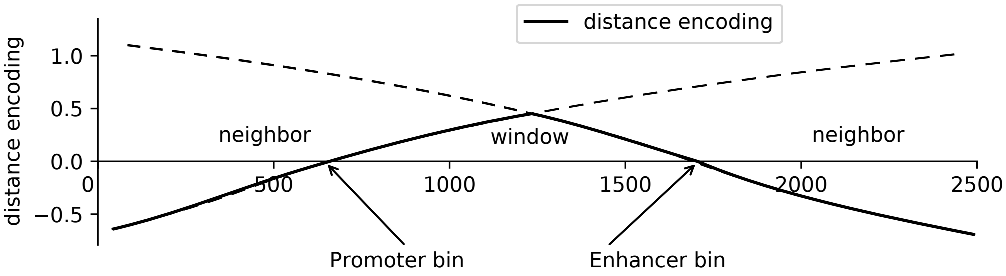

Figure S4: Schematic diagram for distance encoding. The bins in window and neighbor are marked with positive and negative distance encoding, respectively.

### 2 Supplementary Methods

#### 2.1 Preparing features for TrasnEPI

TransEPI compiles the genomic features from large genomic regions of the length of 2.5M base pairs (bp). We partitioned the 2.5Mbp interval into 5000 consecutive bins using the bin size of 500 bp. In each bin, the genomic signals are averaged for each feature, respectively, where the genomic signal is normalized through the arcsinh transformation ($\text{arcsinh}(x) = \ln(x + \sqrt{1 + x^2})$) to reduce the impact of outliers (Ernst and Kellis, 2015; Libbrecht *et al.*, 2019). The genomic features can be classified into 4 groups: (a) CTCF binding sites, (b) DNase I signal, (c) 5 core histone marks (Kundaje *et al.*, 2015), and (d) distance encoding.

1. CTCF binding sites: We downloaded the CTCF ChIP-seq data in narrowPeak format from ENCODE(Davis *et al.*, 2018). The 7th column in narrowPeak file is used as the signal of CTCF binding.

2. DNase-seq signals: We downloaded the DNase I data (P-value track is used because it usually has a high signal-to-noise ratio (Schreiber *et al.*, 2020)) in bigWig format from the Roadmap Epigenomics Project. The Python library pyBigWig (Ramírez *et al.*, 2016) is used to extract the values from bigWig files.

3. Histone modification markers: The same to DNase-seq data.

4. distance encoding: In order to make the model aware of the location of the enhancer and the promoter in each sequence, we encode the distance of each bin to the enhancer or the promoter as an additional feature. The distance encoding is defined as:

$$d = \begin{cases} -\log_2(1 + |\text{index}_i - \text{index}_{\text{a}_1}|), & \text{index}_i < \text{index}_{a_1} \\ \min(\log_2(1 + |\text{index}_i - \text{index}_{a_1}|), \log_2(1 + |\text{index}_i - \text{index}_{a_2}|)), & \text{index}_{a_1} \leq \text{index}_i < \text{index}_{a_2} \\ -\log_2(1 + |\text{index}_i - \text{index}_{\text{a}_2}|), & \text{index}_i > \text{index}_{a_2} \end{cases} \tag{1}$$

where $\text{index}_i, \text{index}_e, \text{index}_p$ are the indexes of the $i$-th bin, the enhancer, and the promoter, respectively. $\text{index}_{a_1} = \min(\text{index}_e, \text{index}_p), \text{index}_{a_2} = \max(\text{index}_e, \text{index}_p)$. A schematic diagram of the distance encoding is shown in Figure S4

#### 2.2 Collecting disease-irrelevant mutations from gnomAD database

The gnomAD database aggregates mutations from the whole genome sequencing data from xxx individuals without diseases. We downloaded the database from `https://gnomad.broadinstitute.org/downloads` (version 2) and randomly sampled $19,715$ single nucleotide variants to compile a disease-irrelevant control dataset. The ratio of the variants on each chromosome and from each type of genomic function region is kept the same to that of GWAS mutations.

#### 2.3 Annotating the genomic function region of mutations

We annotated the genomic function region (e.g.: exon, intron, intergenic region) using ANNOVAR(Wang *et al.*, 2010) against the ENSEMBL gene annotation with two custom options: `--neargene 5000` and `--splicing_threshold 100`.

#### 2.4 Identifying target genes of non-coding mutations

Using TransEPI, we assign a score to each mutation-gene pair which indicates the probability that the genomic loci where the mutation reside in interacts with the promoter of the target gene. We still need a threshold to determine which genes are the targets. Here, we referred to the predictions on the validation dataset and set the threshold to 0.33 because the false-positive rate is below 0.05 when the threshold equals 0.33.

#### 2.5 Ablation study on the contribution of different features

We conducted ablation studies on the genomic features to study their contribution to EPI prediction.

We first studied the contribution of each feature. We excluded a feature from the features each time and re-train the model through the 5-fold cross-validation scheme. As shown in Table S4, we observed that the removal of CTCF induces the most significant decrease in AUC ($-1.89\,\%$) and auPRC ($-5.75\,\%$), followed by DNase I (AUC change = $-1.30\,\%$, auPRC change = $-4.48\,\%$). The removal of a single histone mark has a relatively small impact on the performance. However, when we remove all the histone marks, the decrease in AUC ($-2.38\,\%$) and auPRC ($-7.40\,\%$) surpasses that in the w/o CTCF and the w/o DNase I setting.

Taken together, although CTCF is the most important feature for EPI prediction, the other features, including DNase I signals and some histone marks also greatly contribute to the prediction.
